# Low-coverage reduced representation sequencing reveals subtle within-island genetic structure in Aldabra giant tortoises

**DOI:** 10.1101/2021.11.08.467072

**Authors:** F.G. Çilingir, D. Hansen, N. Bunbury, E. Postma, R. Baxter, L.A. Turnbull, A. Ozgul, C. Grossen

**Affiliations:** Department of Evolutionary Biology and Environmental Studies, University of Zurich, CH-8057 Zurich, Switzerland; Zoological Museum, University of Zurich, CH-8006 Zurich, Switzerland; Indian Ocean Tortoise Alliance, Ile Cerf, Victoria, Republic of Seychelles; Seychelles Islands Foundation, PO Box 853, Victoria, Republic of Seychelles; Centre for Ecology and Conservation, College of Life and Environmental Sciences, University of Exeter, Penryn, Cornwall, TR10 9FE, UK; University of Oxford, Plant Sciences Department, South Parks Road, Oxford, OX1 3RB, UK

## Abstract

*Aldabrachelys gigantea* (Aldabra giant tortoise) is one of only two giant tortoise species left in the world and survives as a single wild population of over 100,000 individuals on Aldabra Atoll, Seychelles. Despite this large current population size, the species faces an uncertain future because of its extremely restricted distribution range and high vulnerability to the projected consequences of climate change. Captive-bred *A*. *gigantea* are increasingly used in rewilding programs across the region, where they are introduced to replace extinct giant tortoises in an attempt to functionally resurrect degraded island ecosystems. However, there has been little consideration of the current levels of genetic variation and differentiation within and among the islands on Aldabra. As previous microsatellite studies were inconclusive, we combined low-coverage and double digest restriction associated DNA (ddRAD) sequencing to analyze samples from 33 tortoises (11 from each main island). Using 5,426 variant sites within the tortoise genome, we detected patterns of population structure *within* two of the three studied islands, but no differentiation between the islands. These unexpected results highlight the importance of using genome-wide genetic markers to capture higher-resolution genetic structure to inform future management plans, even in a seemingly panmictic population. We show that low-coverage ddRAD sequencing provides an affordable alternative approach to conservation genomic projects of non-model species with large genomes.

## Introduction

Many endangered species are restricted to a single or a small number of remnant populations. Management efforts often include introductions from these source populations to other suitable locations to lessen the risk of extinction or because the species in question are ecosystem engineers and can be used to restore degraded habitats elsewhere. However, such interventions have important implications for the genetic future of the newly founded population. As only a subset of the individuals in the source population can be moved, genetic diversity is at risk to be lost and artificial population structure may be created in the new populations. Genetic diversity is essential for the adaptive potential of a species, particularly in the face of environmental changes and disease outbreaks (Reed, 2005; Reed & Frankham, 2003). Hence, management decisions need to be carefully planned to take the genetic characteristics of the source populations into account to aim at retaining as much genetic diversity as possible (Hoban et al., 2021).

One problem with assessing current genetic characteristics of endangered non-model species is that suitable marker systems, such as simple sets of microsatellites, are often unavailable. Next-generation sequencing provides promising tools at decreasing costs (Davey & Blaxter, 2010; Hayden, 2014). However, it can still be financially overwhelming and (if not outsourced) bioinformatically challenging to generate high-quality whole genomes, especially for species with large genomes, and because more than a handful of sequenced individuals are needed for population genomics studies (Corlett, 2017; Shafer et al., 2015). One potential solution is to use reduced representation sequencing, such as restriction-associated DNA (RAD) sequencing, which does not require a reference genome and is generally cost-effective (Andrews, Good, Miller, Luikart, & Hohenlohe, 2016; Davey & Blaxter, 2010). Financial and computational costs of whole-genome sequencing of many individuals can be further reduced by adopting a low-depth sequencing strategy (Pasaniuc et al., 2012), where information on the whole genome is obtained, but at low coverage (generally 1–2x). This approach risks loss of genotype accuracy, which can be overcome by inferring genotype likelihoods (Fumagalli, Vieira, Linderoth, & Nielsen, 2014; Korneliussen, Albrechtsen, & Nielsen, 2014). This genotype-free estimation of allele frequencies has been shown to reduce biases and improve demographic inference from RAD-seq data (Warmuth & Ellegren, 2019). Interestingly, to date, only a small number of studies have combined RAD and genotype-free estimation of allele frequency estimation approaches (e.g. Bay, Taylor, & Schluter, 2019; Breusing, Johnson, Vrijenhoek, & Young, 2019; Peart et al., 2020; Záveská et al., 2019).

Here, we use low-coverage ddRAD sequencing as a time- and cost-effective approach for the population genetic analysis of *Aldabrachelys gigantea*, Schweigger 1812 (Aldabra giant tortoise), a flagship and keystone species lacking both a suitable marker set and a reference genome. *A*. *gigantea* is one of only two giant tortoise species left in the world together with the Galápagos giant tortoise complex, *Chelonoidis niger*, (Turtle Taxonomy Working Group, 2017) and is endemic to Aldabra Atoll, Seychelles. The species is currently listed as Vulnerable by the IUCN Red List (version 2.3), due to its limited distribution in the wild and threats posed by climate change. It is the only survivor of at least nine endemic giant tortoise species that once lived on Western Indian Ocean islands (Austin, Nicholas Arnold, & Bour, 2003; Palkovacs, Gerlach, & Caccone, 2002) and occupies a prominent functional role in shaping and sustaining large-scale vegetation dynamics as it is the largest frugivore and herbivore in its island ecosystem (Hansen, 2015; Hnatiuk, Woodell, & Bourn, 1976; Merton, Bourn, & Hnatiuk, 1976). Therefore, *A*. *gigantea* are currently used to help restore degraded native ecosystems on several other Western Indian Ocean Islands (Griffiths, Hansen, Jones, Zuël, & Harris, 2011; Griffiths, Zuël, Jones, Ahamud, & Harris, 2013; Hansen, Josh Donlan, Griffiths, & Campbell, 2010).

*A*. *gigantea* was on the verge of extinction in the late 19th century due to excessive harvesting, with a population low in around 1870 of somewhere between < 1000 and a few thousand tortoises (Bourn et al., 1999; Stoddart & Peake, 1979). Thanks to calls for protection from Charles Darwin and others in 1874, the number of *A*. *gigantea* increased quickly to several tens of thousands in the 1960s to today’s stable population of well over 100,000 individuals (Turnbull et al., 2015). The Aldabra population is divided into several subpopulations across the different islands that make up the atoll.

Here, we provide new sampling and analysis that acts as a case study for the conservation genetic analysis of a non-model species using low coverage sequencing combined with double-digest restriction site-associated DNA sequencing (ddRADseq, Peterson, Weber, Kay, Fisher, & Hoekstra, 2012). Our specific aims are:

1. To quantify the overall genetic structure of the endemic *A*. *gigantea* population
2. To determine whether there are significant differences in the genetic composition of the species among and within islands.

## Materials and Methods

### Study system

The endemic distribution of *A*. *gigantea* is restricted to Aldabra Atoll, in the southern Seychelles. The atoll consists of four main islands, Grande Terre, Malabar, Polymnie, and Picard (Fig. 1A), separated by channels and enclosing a shallow lagoon. On the atoll, giant tortoises are unevenly distributed across the three largest islands (Polymnie, the smallest main island, has no tortoises) due to environmental differences (e.g. terrain, food, freshwater resources, shade availability) and differences in exploitation history (Bourn & Coe, 1978; Turnbull et al., 2015; Walton et al., 2019). Effective conservation management measures saving the species from extinction in the late 19th century included the reintroduction of tortoises to Picard and atoll-wide invasive species control (Bourn et al., 1999; Bunbury et al., 2018; Stoddart & Peake, 1979; Turnbull et al., 2015). The largest population lives on Grande Terre, with the second-largest on Malabar. Polymnie, surrounded by deep channels, remains empty of *A*. *gigantea*, while Picard has been re-populated in several translocations of tortoises from Malabar and Grande Terre since the early 1900s, with the last occurring in the 1980s. An unknown number of *A*. *gigantea* occur around the globe in captivity, semi-natural or re-wilded populations (Hansen, Donlan, Griffiths, & Campbell, 2010).

**Fig. 1.**
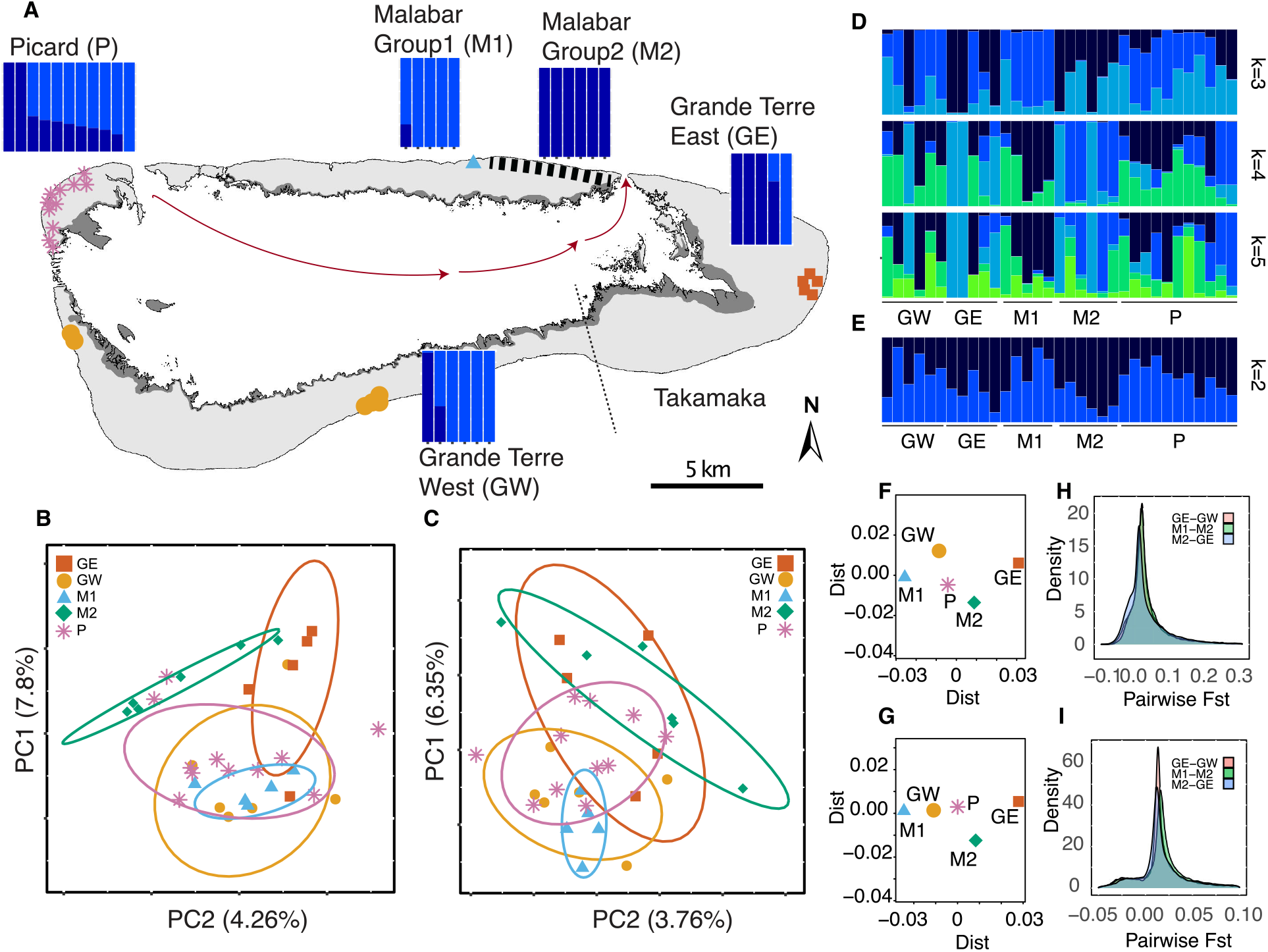
**A)** Aldabra Atoll’s four main islands. The curved arrow within the map indicates the direction of the ocean currents. Gray shaded areas show the mangrove distribution within the atoll. Dashed lines show the region of Takamaka. Every colored mark on the map represents a sampled tortoise. The color and shape of the marks indicate a distinct sampling location within each island. The area delimited by the serrated line shows approximate sampling locations of the rest of the Malabar samples on the north-eastern side of the island. Each bar above the islands corresponds to one individual sampled there and shows its admixture proportions estimated with the main dataset assuming two ancestral populations (k=2). Light blue bars: cluster A, Dark blue bars: cluster B. **B)** Five genetic clusters are shown on the PCA plot of the main dataset and **C)** the downsampled dataset. Every colored mark represents an individual. Malabar Group 2 individuals are shown with green diamonds. **D)** Admixture proportions of all the individuals estimated with the main dataset assuming k=3-5, **E)** with downsampled dataset assuming k=2. **F)** MDS of the pairwise Fst values estimated for each group with the main dataset and **G)** the downsampled dataset, each mark represents the whole group. **H)** Density plot of the sliding window analysis of pairwise Fst between three genetic groups representing within- and among-island genetic differentiation, estimated with the main dataset, (GE-GW and M1-M2, within Grande Terre and Malabar, respectively; GE-M2, among Grande Terre and Malabar), **I)** estimated with the downsampled dataset.

### Sample collection and DNA extraction

In 2012 and 2013, approximately 100μl of blood were drawn from the cephalic vein of an extended front limb of 33 adult *A*. *gigantea* individuals representing the three main islands of Aldabra, which are inhabited by tortoises: 11 from Picard, 11 from eastern Malabar, and 11 from Grande Terre (West, n=6; East, n=5) (Fig. 1A, Suppl. Table 1). Absolute ethanol was added to the blood samples in a 1:20 ratio to prevent coagulation (Wietlisbach, 2017). All samples were stored at room temperature until arrival in the lab and then at −80°C until DNA extraction.

DNA extraction was performed with 3μl of blood (in ethanol) per sample, using the sbeadex™ kit (LGC Genomics, Middlesex, UK), following the manufacturer’s protocol for DNA extraction from nucleated red blood cells. Genomic DNA concentrations were measured with a dsDNA Broad Range Assay Kit (Qubit 2.0 Fluorometer, Invitrogen, Carlsbad).

### ddRAD-seq library preparation and sequencing

To keep sequencing costs as low as possible, we used a reduced-representation genome sequencing approach, specifically the double-digest restriction site-associated DNA sequencing (ddRAD-seq, Peterson et al., 2012). Restriction enzymes were selected based on *in silico* double digest runs, using the SimRAD package within R v4.0.3 (Lepais & Weir, 2014; R Core Team, 2020). Enzyme combinations of EcorI-TaqI, EcoRI-MspI, and EcoRI-BfaI were tested using *in silico* restriction digests, performed on the basis of a *Chelonoidis abingdonii* (Galápagos giant tortoise) genome, which is the phylogenetically closest available genome for *A*. *gigantea* (ca. 35-40 My of divergence time; (Kehlmaier et al., 2019; Quesada et al., 2019). We aimed for approximately 50,000 *in silico* RAD loci, which was achieved with the selected enzyme combination EcoRI-BfaI with a target size selection window of 300–350 bp (52’000 expected ddRAD loci, Suppl. Fig. 1).

We used 100 ng genomic DNA from each sample (n=33) for the digestion. A single ddRAD-seq library was prepared by processing the 33 samples following the protocol by Peterson et al. (2012) with slight modifications as described in (Çilingir, Hansen, Ozgul, & Grossen, 2021). Two additional samples representing captive populations (one from Mauritius, one from Rodrigues) were also processed and sequenced, but not included in this study, which was focused on the wild populations. Briefly, after double digestion, the products were cleaned with a 1.0x ratio of AMPure XP beads. Next, the P1 adapters containing the inline barcodes unique to each sample (Peterson et al., 2012), and with an EcoRI overhang and the P2 adapter with a BfaI overhang were ligated to the restricted DNA. Then, equal amounts of individually barcoded DNA were pooled. The double size selection was performed with a total of 300ul pooled aliquot by treatment with 0.5x and 0.12x AMPure XP beads. After the size selection, 8 PCR cycles were run using the common PCR1 and the PCR2 primers, which include a standard Illumina index (Peterson et al., 2012). In our case only one index was used as there was only one sequencing library prepared). A final AMPure XP beads clean up followed with a 0.6x bead ratio. The quality check of the final library fragment size range was performed with a Bioanalyzer High Sensitivity DNA kit (Agilent, Santa Clara, CA). Finally, 10 picomoles of the quality-checked library were sequenced on an Illumina Miseq platform for a paired-end run on one lane at the Genetic Diversity Center, ETH Zurich, Switzerland, yielding read lengths of 300 bp.

Data quality of the sequences was assessed using FastQC v0.11.9 (S. Andrews, 2010), and the adaptor sequences of the Illumina sequencing platform were trimmed using Trimmomatic v0.39 (Bolger, Lohse, & Usadel, 2014) (ILLUMINACLIP:2:30:10:2). Adapter-trimmed data were demultiplexed in Stacks v2.53 (Rochette, Rivera-Colón, & Catchen, 2019) using process_radtags and allowing one barcode mismatch. At this step also all reads containing at least one N (uncalled base) were removed. Quality filtering of the demultiplexed data was done with Trimmomatic (Bolger et al., 2014) requiring an average Phred quality score per entire read of at least 20 (AVGQUAL:20), an average quality of 10 in a sliding window of 30 before cutting the read (SLIDINGWINDOW:30:10), bases were cut off the end of the read if the quality dropped below 19 (TRAILING:19) and the first 10 bases were cropped to remove the enzyme cut sequence (HEADCROP:10).

### Alignment to a reference genome, estimation of sequencing depth, and downsampling

After quality filtering, the paired reads were aligned to the *C*. *abingdonii* reference genome (NCBI BioProject PRJNA611832) using BWA-MEM version 0.7.17 (Li & Durbin, 2009). Calculation of the average per site sequencing depth for each individual was done in three following steps. First, SAMtools (Li et al., 2009) was used to extract properly paired reads with mapping quality of >20 from the BAM file of individual GrdTr_11 (the individual with the highest number of sequence reads, Suppl. Table 1). Next, for each individual, all positions with at least one read were retained within a bed file by using bedtools v2.29.2 (Quinlan & Hall, 2010). Subsequently, per-site sequencing depth per individual was calculated using SAMtools (Li et al., 2009) based on the range given by the bed file (all sites with at least 1x coverage).

Because the average sequencing depth per individual varied considerably, we repeated the major analyses after downsampling the forward and reverse Fastq files of each sample to equalize the number of reads per individual with seqtk v1.3 (https://github.com/lh3/seqtk) to 154,599 reads, (number of reads of individual Picard_2, third-lowest read count, Suppl. Table 1).

### Estimation of genotype likelihoods

As the mean sequence coverage per sample was low (2.28x; range: 0.2–6.1x, Suppl. Table 1), the uncertainty of genotypes was accounted for in the subsequent analyses by computing the genotype likelihoods at variant sites instead of calling genotypes. Accordingly, the read alignments of all 33 individuals were processed with ANGSD v0.93 (Korneliussen et al., 2014), a software developed for genomic analyses of low coverage data. The GATK model was used (McKenna et al., 2010), and major and minor alleles were directly inferred from the genotype likelihoods (doMajorMinor 1, doMaf 1). Quality filtering for the subsequent downstream analyses was performed as follows: Only properly paired (only_proper_pairs 1) and unique reads (uniquieOnly 1) were used, and only biallelic sites were retained (skipTrialleleic 1). Nucleotides with base qualities lower than 20 were discarded. Excess of SNPs around indels and excessive mismatches with the reference were corrected by realignment (C50, baq 1, Li, 2011). Reads with a mapping quality lower than 20 were discarded.

Additionally, for the estimation of genotype likelihoods, only SNPs with a p-value < 10^−6 (the significance threshold for polymorphism detection) and heterozygosity < 0.5 were retained, the latter to exclude potential paralogs (Hardy, 1908; Hohenlohe, Amish, Catchen, Allendorf, & Luikart, 2011). Further filters were applied depending on the analysis. For the population genetic structure analyses, sites with read data in fewer than 30 of the 33 samples were excluded (minimum representation among samples > 90%, -;minInd 30). The minimum depth of sites to be retained was also set to 30, and hence, on average, at least one read per individual was required. The maximum depth per site was set as the sum of the average sequencing depth and two times the standard deviation (373 for the main dataset, 128 for the downsampled dataset). For the estimation of genetic differentiation and diversity, which were calculated per group, at least 50% of the samples in a particular group had to be represented (minInd = 50% of all individuals in a group). The minimum depth for each group was set to the minimum number of individuals allowed (50% of the overall individuals within a group) and the maximum depth was the average plus two times the standard deviation for each group.

### Population genetic structure

For a first overview of the population structure, a principal component analysis was carried out with PCAngsd v09.85 (Meisner & Albrechtsen, 2018) with an additional minor allele frequency (MAF) filter of 0.01 or 0.05. As a complementary population structure analysis, we used the clustering tool NGSAdmix (Meisner & Albrechtsen, 2018; Skotte, Korneliussen, & Albrechtsen, 2013). Similar to the Bayesian clustering method STRUCTURE (Pritchard, Stephens, & Donnelly, 2000), NGSAdmix allows the estimation of individual admixture proportions by assigning individuals to different clusters. While a PCA allows the assumption-free visualization of the genetic relatedness among individuals, NGSAdmix tries to minimize the within-group variation to define genetic groups and estimate individual admixture proportions (Meisner & Albrechtsen, 2018; Skotte et al., 2013). To use NGSAdmix, it is recommended to perform LD pruning (i.e. to filter sites based on pairwise linkage disequilibria) as the program assumes the independence of genomic loci (Skotte et al. 2013). Hence, pairwise linkage disequilibria (LD) were calculated using ngsLD (Fox, Wright, Fumagalli, & Vieira, 2019) and LD pruning was performed by allowing a maximum among SNP distance of 100 kilobases and a minimum weight of 0.5. A total of 100 replicates were performed for each NGSAdmix run and the number of clusters (k) varied between two and 10. The results were analyzed and visualized with CLUMPAK (Kopelman, Mayzel, Jakobsson, Rosenberg, & Mayrose, 2015), and the log-likelihoods calculated for each run were visualized in R (R Core Team, 2020).

### Estimation of genetic differentiation and diversity comparison

As a measure of population differentiation, weighted pairwise Fst values were calculated between each population of three different islands (Picard, Malabar, and Grande Terre) and each general sampling location within the islands (total of five locations, Fig. 1) by using ANGSD (Korneliussen et al., 2014) and realSFS (a module of ANGSD). For each population/group, the site allele frequency (SAF) likelihoods were estimated based on individual genotype likelihoods (see section Estimation of genotype likelihoods) with the - doSAF 1 option of ANGSD (Korneliussen et al., 2014). The SAF was polarised with the reference genome as no ancestral sequences were available. Then, folded Site Frequency Spectra (SFS) were calculated for each population and Fst metrics were estimated using 2D-SFS and the option -whichFst 1. To visualize the genetic differentiation between all groups/populations, multidimensional scaling (MDS) was applied to the pairwise Fst matrix using the cmdscale function in R (R Core Team, 2020). Moreover, a heatmap of the pairwise Fst values was generated with ggplot2 (Wickham, 2016) in R (R Core Team, 2020). Additionally, to account for potential local effects along the genome, a sliding window analysis of the pairwise Fsts was performed for the comparison within Malabar and Grande Terre Islands, as well as among Grande Terre and Malabar with a window and step size of 50 kilobases (non-overlapping windows, excluding windows with < 10 sites).

Possible differences of genetic diversity among the five groups defined above (see also Fig. 1) were investigated by calculating average number of pairwise differences or nucleotide diversity (π; (Tajima, 1989)) and population mutation rate (Watterson’s θ; (Watterson, 1975)). Both measures were based on SFS estimates and performed with the realSFS and ThetaStat modules in ANGSD (Korneliussen et al., 2014). Estimates of Watterson’s θ and π were obtained per genome region via a sliding window analysis with a window and step size of 10 kilobases (non-overlapping windows, excluding windows with < 10 sites). A Tukey’s range test (David & Tukey, 1977) was applied to compare the diversity measures among different groups. Since Tukey’s range test is a post-hoc test, initially ANOVA was performed on the data.

## Results

### Genotype likelihood analysis

The sequencing effort yielded 24,942,560 raw reads for a total of 35 samples. After adapter removal, quality checking, and demultiplexing, an average of 1,188,685 (range: 129,188– 2,610,656, see also Suppl. Table 1) reads per sample were retained in the main dataset. The mean mapping rate was 96.5% (range: 94.7–97.0%, Suppl. Table 1) resulting in a mean sequencing depth per sample of 2.28x (range: 0.2–6.1x, Suppl. Table 1). In the downsampled dataset, all individual fastq files were downsampled to 154,599 sequencing reads and the mean sequencing depth per sample was calculated as 0.56x (range: 0.2–0.8x). The genotype likelihood analysis with ANGSD (Korneliussen et al., 2014) resulted in 238,995,840 sites, 6,153 of which were retained as variant sites (SNPs). MAF filtering for > 0.05 yielded 5,426 SNPs. 189,369,839 sites were obtained using the downsampled dataset, 1755 of which were retained as variant sites (SNPs). MAF filtering for > 0.05 yielded 1,632 SNPs.

### Population genetic structure

We were primarily interested in the overall genetic structure and differentiation among islands. The PCA of the main dataset revealed two distinct clusters (PC1:7.8% and PC2: 4.26%, Fig. 1B). One cluster was represented by five of the six individuals from Grande Terre West (GW), five individuals from Malabar (now termed ‘Malabar Group 1’ or M1), one Grande Terre East (GE) individual, and the majority of all Picard (P) individuals (n=9). The second cluster grouped four of the five GE individuals, six Malabar individuals (now termed ‘Malabar Group 2’ or M2), one GW individual, and the two remaining P individuals (Fig. 1B). M1 was clearly separated from M2, and GW and GE formed overlapping but differing groups. P individuals overlapped with all four groups. The PCA of the downsampled dataset (PC1: 6.35% and PC2:3.76% Fig. 1C) resulted in a less clear resolution but confirmed the general population structure described above for the full data set. Changing the minor allele cut-off (MAF > 0.01 vs MAF > 0.05) had no effect on the PCA structure (Suppl. Fig. 2A & 2B).

Next, we wanted to investigate if the observed population structure is consistent with two genetic groups as indicated by the PCA and we aimed at estimating admixture proportions. For this analysis, a total of 3781 LD-pruned SNPs with MAF > 0.05 were used. The admixture proportions indicated that when the number of putative clusters was assumed to be two (k=2, was the most likely number of k based on log-likelihoods, Suppl. Fig. 3A), the M1 individuals were assigned to cluster A and the M2 individuals were assigned to cluster B, except for one individual of M1, which showed mixed ancestry (Fig. 1A). Also, Grande Terre showed high within-island differentiation with most of GE assigned to cluster B and most of GW to cluster A. All P individuals except for two assigned to cluster B showed mixed ancestry. Hence, genetic groups of different islands were assigned to the same clusters (M1 with GW, and M2 with GE). This high within-island differentiation on Malabar and Grande Terre but lower among-island differentiation confirmed the outcome from the PCA (Fig. 1B). Under a scenario of 3–10 hypothetical clusters, all groups showed mixed ancestry (Fig. 1D, Suppl. Fig. 4). The results of the admixture analysis based on the downsampled dataset including a total of 1120 LD-pruned SNPs with MAF > 0.05 were consistent with the results based on the main dataset (Fig. 1E; Suppl. Fig. 3B, 5 & 6). Admixture proportions obtained with the two datasets were positively correlated, but the retained resolution was considerably lower (Fig. 1E, Suppl. Fig. 6).

### Estimation of genetic differentiation and summary statistics

All pairwise Fst estimates calculated among the three Aldabra Islands in the study (Picard, Malabar, and Grande Terre) were zero, suggesting no evidence for among-island differentiation. As expected from the PCA and the admixture proportion analysis, the major differentiation was found between M1 and GE (0.06), followed by GE and W (0.041), and M1 and 2 (0.039) (Fig. 2, Fig. 1F). To account for possible local effects along the genome, we also compared within- and among-island differentiation of Grande Terre and Malabar by performing a sliding window analysis of pairwise Fsts. The analysis confirmed a slightly lower among-island differentiation between Malabar and Grande Terre than within-island differentiation on Malabar. The analysis of the main dataset indicated a level of within-island differentiation on Grande Terre similar to that on Malabar, but this differentiation was lower when analyzing the downsampled data set (Fig. 1H, 1I). The pairwise Fst estimations with the downsampled dataset confirmed the major finding of within-island differentiation (Suppl. Table 2, Fig. 1).

**Fig. 2.**
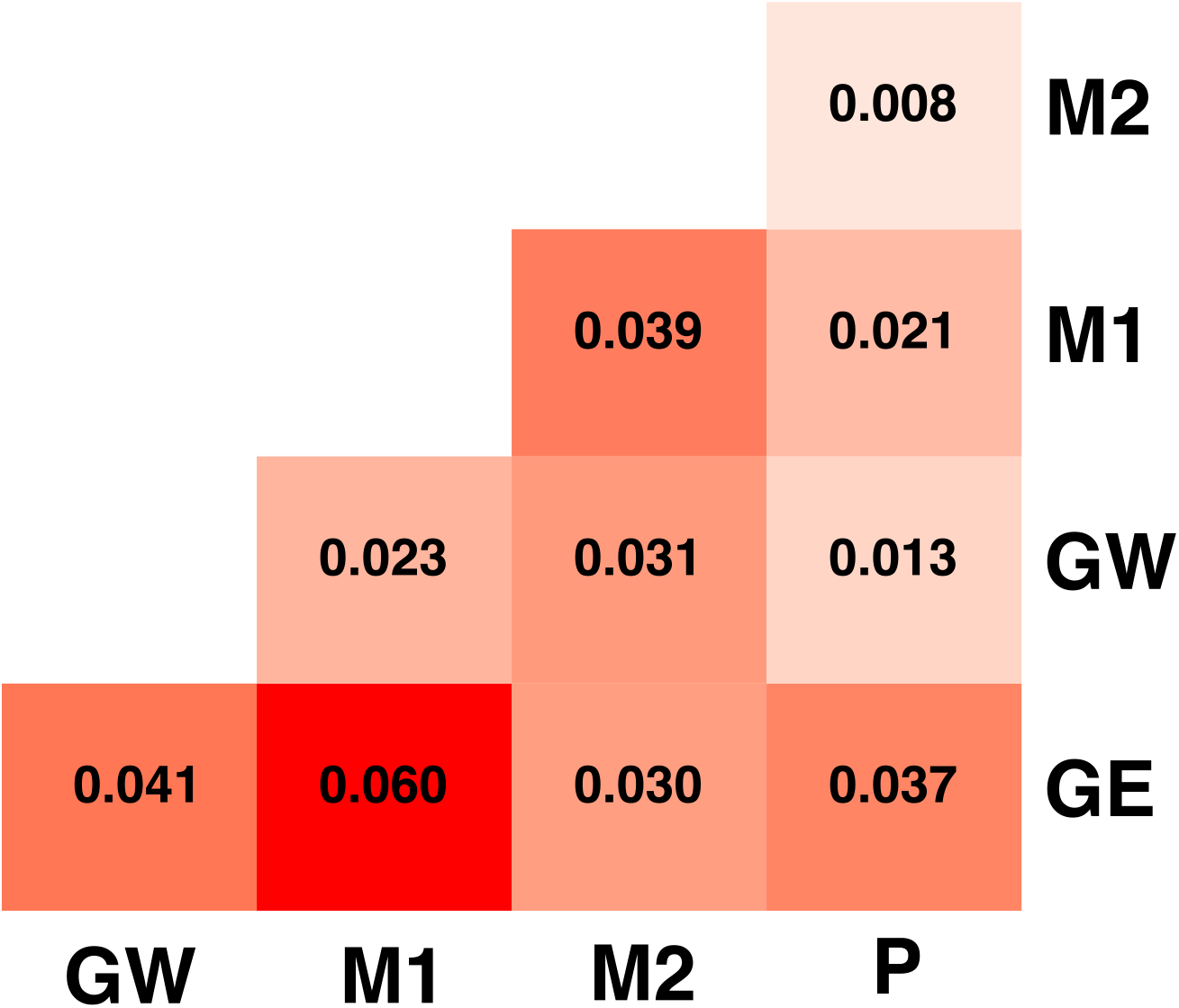
Heatmap of pairwise genetic differentiation (measured as Fst), estimated for five different locations using the main dataset

Mean Watterson’s θ values of all the groups ranged from 0.00139 to 0.00167, with P having the highest estimate, and GE the lowest (Fig. 3A) suggesting highest genetic diversity in P. There was significant variation among the groups, F (4, 596387)=459, p < 2e-16. All the groups’ mean Watterson’s θ values were significantly different from each other at p < 0.05. Mean π per group was 0.00143–0.00153, with P having the highest and M2 the lowest values (Fig. 3B), again suggesting highest genetic diversity in P. Although the absolute differences among groups were small, there was significant variation among the groups, F (4, 596387)=75.23, p < 2e-16. Mean π values of all the groups differed from each other, except for GE, which didn’t differ from M2 nor GW. While both Wattersons θ and π as well the analyses with the downsampled dataset suggested highest diversity in P, differences in diversity among the other groups were small (but significant) and the relative order of groups differed between analyses.

**Fig. 3.**
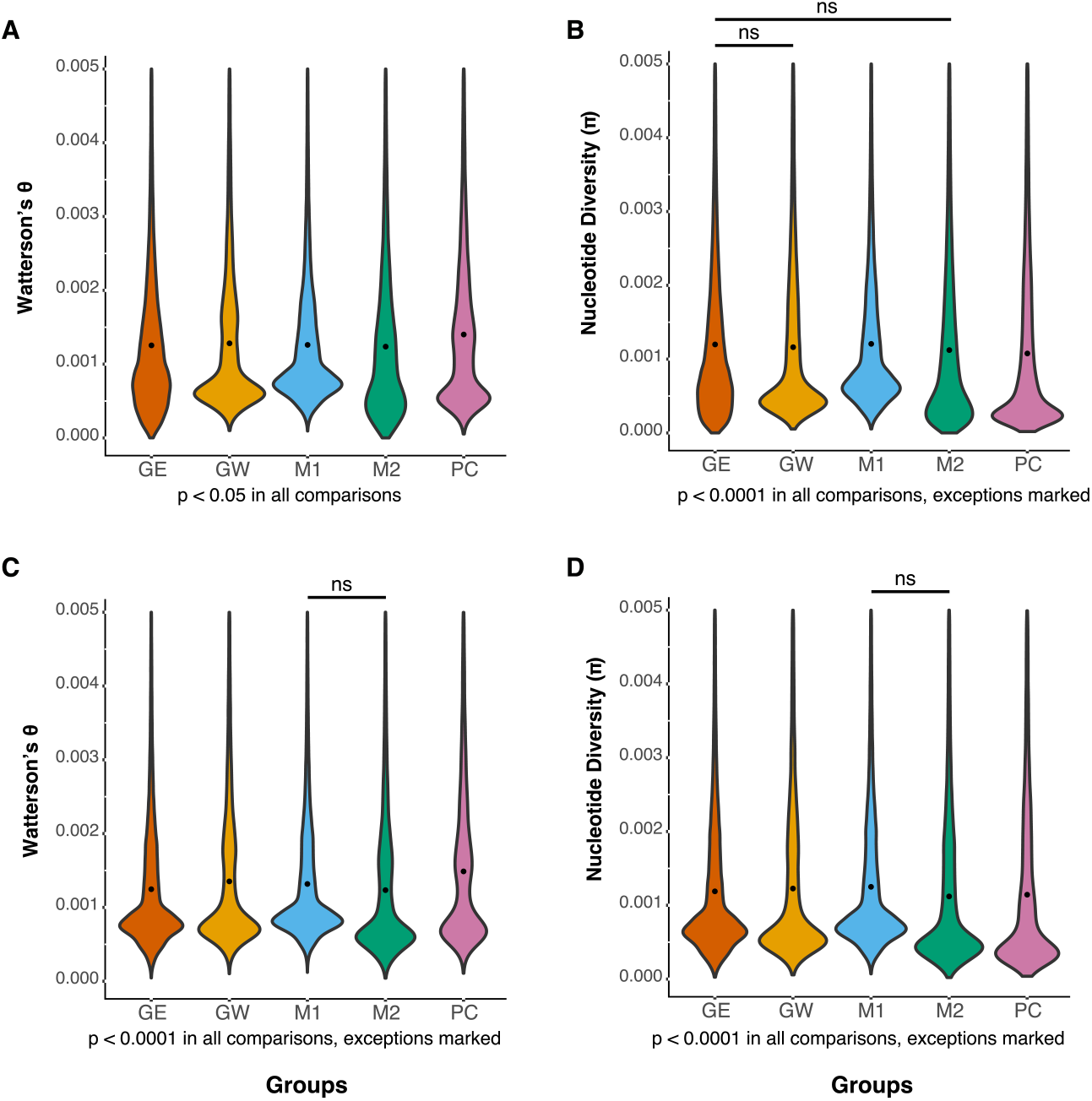
**A)** Per-site estimates of Watterson’s θ **B)** and nucleotide diversity (π) obtained via a sliding window analysis performed with the main dataset; **C)** per-site estimates of Watterson’s θ **D)** and nucleotide diversity (π) obtained via a sliding window analysis performed with the downsampled dataset. Each group is colored the same as in Fig. 1 (orange, Grande Terre East; yellow, Grande Terre West; blue, Malabar Group 1; green Malabar Group 2; pink, Picard) and the average value per each group is indicated with a black dot.

## Discussion

We used low-coverage RAD sequencing to investigate the population genetic structure and variation of the endemic *A*. *gigantea* population. Our data, although relying on a relatively small sample size (5-11 per island/sampling location), not only revealed the subtle genetic structure of previously bottlenecked populations but suggest a potentially greater role of passive movement between islands via water in a terrestrial species than previously expected.

Our study is one of few to focus on a combination of reduced representation sequencing and the genotype likelihood approach to study the population genomics of a non-commercial and non-model species. Our case study supports the use of low-coverage ddRAD sequencing instead of the low coverage whole-genome sequencing (Lou, Jacobs, Wilder, & Therkildsen, 2021), which is still costly for large genomes and/or sample sizes. Sequencing costs depend on the platform, but could be as low as 5.25 USD per sample using our approach (2-3x coverage or 0.3Gb). In contrast, a low-coverage whole genome resequencing project for a genome of about 2.4 Gb (the approximate genome size of *A*. *gigantea*) would result in sequencing costs of about 105 USD per sample (2-3x coverage or 6Gb). Our reduced representation approach could therefore be particularly useful for species with very large genomes.

We also investigated the effects of unevenly distributed depth of sequencing per individual by repeating all analyses with a downsampled dataset. We showed that the results obtained with both datasets were consistent, but the downsampling led to a loss of resolution, especially for the admixture analysis. The smaller number of loci and among-locus variation in coverage known for RAD (Davey et al., 2013; O’Leary, Puritz, Willis, Hollenbeck, & Portnoy, 2018) may mean that there is a minimum acceptable depth of coverage for this technique.

### Unexpected partitioning of genetic structure

We found lower among-island than within-island differentiation. Specifically, we found two main clusters of genetic variation (Fig. 1A): M2, and all but one individual from GE represented an eastern group. M1 and all but one individual from GW represented a western group. The P samples were assigned to both clusters, which was expected, given that the original population of Picard was extirpated in the 1800s, and the current population originates from re-introduced tortoises from Grande Terre and Malabar (Bourn et al., 1999). These findings were supported by the PCA and the pairwise Fst analyses, which showed minor differentiation between Picard and the other islands, but relatively strong differentiation within Grande Terre and Malabar. The genetic differentiation between GE and GW suggests that connectivity along the east-west axis of the island may be limited. This is in agreement with behavioral, ecological and geographic observations (Bourn & Coe, 1978; Gibson & Hamilton, 1984; Swingland, North, Dennis, & Parker, 1989), and a previous study by Balmer et al. (2011). Areas of thick *Pemphis* scrub and deeply fissured rocks appear to limit the movement of tortoises (Gibson & Hamilton, 1984). Hence, geographical barriers such as the region around Takamaka (dotted line in Fig. 1A) that include deeply fissured limestone and thick *Pemphis* scrub, probably explain the observed substructure on Grande Terre (Balmer et al., 2011; Bourn & Coe, 1978).

More surprising was the low differentiation between M2 and GE. Occasional movement of tortoises carried by tidal currents from the mangrove area in Grande Terre East to the coastal area of M2 may cause inter-island gene flow (Fig. 1A). The tortoises often spend days or weeks in the muddy mangroves of Grande Terre East. Sometimes they move against tidal waters rushing out or in, with a risk of being swept away and tortoises can even be spotted adrift in the open ocean outside the reef (Hansen et al., 2017). Ocean currents are increasingly acknowledged for their importance in shaping population structure (Arjona et al., 2020; White et al., 2010), also for terrestrial reptiles (Calsbeek & Smith, 2003; Hawlitschek, Ramírez Garrido, & Glaw, 2017). The movement of animals by humans could also explain the low differentiation. Although it is known that animals were transported from Grande Terre and Malabar to Picard for conservation purposes, there is no record of animals being transported from Grande Terre to Malabar or vice versa. It is therefore reasonable to assume that a direct route was taken for the tortoises en route to Picard, given that managing/transporting giant tortoises is considerable effort. Eventually, more samples from both of these populations, as well as outgroups to quantify the magnitude of the flow and try to date it, are needed to confirm our hypothesis of across-water gene flow. Evidence for ongoing gene flow over tens of generations would support our current hypothesis. The higher within-island differentiation between M1 and M2 could be a result of the aforementioned flow to Malabar, but limited gene flow with M2. The vegetation between the regions is very dense pemphis with the exception of the coastal path and a previous study on tortoise habitat use showed that Malabar tortoises in general have smaller home ranges compared to residents of other islands (Walton et al., 2019). Perhaps, newly arrived Malabar residents do not move as far as their conspecifics on Grande Terre and Picard, and eventually gene flow is limited between the two groups within Malabar.

One potential reason for a lack of genetic differentiation between the GW and P is the movement of P tortoises to GW via the wide channel and islets between the two islands (Fig. 1A), so that admixed P individuals influenced the GW group. However, this genetic similarity is likely to be driven by the founding history of the Picard population, which received individuals from Grande Terre (Bourn et al., 1999).

In summary, our findings suggest a subtle and unexpected signal of east-west population structure in *A*. *gigantea*, mainly correlated with landscape features as well as human-induced reintroductions (primarily on Picard). Seawater may play a less important role as a barrier than has been previously assumed (Balmer et al., 2011; Grubb, 1971), instead water currents may support movements. Balmer et al. (2011) found no variation at the mitochondrial control region and there is currently no evidence for an ancient split between the two genetic groups. Given the very long generation time of giant tortoises, the substructure could still be several hundred years old and predate the species bottleneck.

### Conservation and research implications

Our study has several conservation implications. Our findings suggest that if giant tortoises from Aldabra are to be used for translocations, translocated individuals should ideally be equally distributed from the east and west axes of Malabar and/or Grande Terre to represent the overall genetic variation in the wild population. Individuals from Picard were assigned to both genetic groups, showed the highest genetic diversity and the island hosts the research station. Hence, individuals taken only from Picard could be a valuable alternative and logistically more feasible. In any case, it is advised to translocate as many individuals as possible to minimize founder effects (Frankham, Ballou, Briscoe, & Ballou, 2007). This should facilitate genetic management and monitoring of ongoing and future rewilding projects, including spatially larger projects in Madagascar (Pedrono et al., 2013), to maximize the evolutionary potential and survival of rewilded populations. As previously suggested by Balmer et al. (2011), we found no evidence for large differences in genetic diversity among the main islands and there is therefore no need for translocations between islands. The similar diversity among islands also suggests that the observed population structure is unlikely to be explained by the species bottleneck, for instance by much stronger reduction and then isolation of the eastern group. However, we caution that our method of low-coverage ddRAD has not been tested sufficiently for its reliability on the estimation of exact diversity measures (see recommendations from Lou et al., 2021).

In conclusion, our study underlines the importance of genetically informed management decisions by showing unexpected population structure as previously discovered in Iberian wolves, Peruvian diving-petrels, and Atlantic puffins, among others (Cristofari et al., 2019; Kersten et al., 2021; Silva et al., 2018). It most notably shows that land- and seascape genetics should go hand in hand because terrestrial organisms living close to the sea could be influenced by both. Finally, considering that both reduced representation sequencing and low-coverage approaches aim to decrease costs, our approach could be used in the population genomics of other vertebrates to address similar research questions.

## Data Accessibility

Data for this study are uploaded at the NCBI Sequence Read Archive with accession numbers SRR14611971-SRR14612003.

## Acknowledgments

We would like to thank the Seychelles Islands Foundation and Aldabra research staff for their invaluable help in the field and their support in exporting the blood samples. The former Zurich-Aldabra Research Platform was involved in the blood sampling, and we especially thank Gabriela Schaepman-Strub for her support and contributions. We thank Xenia Wietlisbach for her help with sample management. We would also like to thank Claudia Michel for her contributions to the optimization of the ddRAD-seq protocol. All the wet-lab work and the sequencing efforts required for this study were conducted in the Genetic Diversity Center, ETH Zürich. During the course of this study, F. Gözde Çilingir was funded by the Swiss Government Excellence Scholarship for Postdocs and the Research Funding from the Research Talent Development Fund of the University of Zürich. The Georges and Antoine Claraz Foundation contributed to the sequencing costs of this project.

## Literature Cited

Andrews, K. R., Good, J. M., Miller, M. R., Luikart, G., & Hohenlohe, P. A. (2016). Harnessing the power of RADseq for ecological and evolutionary genomics. Nature Reviews Genetics, 17(2), 81–92.

Andrews, S. (2010). FastQC: A Quality Control Tool for High Throughput Sequence Data. Retrieved from http://www.bioinformatics.babraham.ac.uk/projects/fastqc/

Arjona, Y., Fernández-López, J., Navascués, M., Alvarez, N., Nogales, M., & Vargas, P. (2020). Linking seascape with landscape genetics: Oceanic currents favour colonization across the Galápagos Islands by a coastal plant. Journal of Biogeography, 47(12), 2622–2633.

Austin, J. J., Nicholas Arnold, E., & Bour, R. (2003). Was there a second adaptive radiation of giant tortoises in the Indian Ocean? Using mitochondrial DNA to investigate speciation and biogeography of Aldabrachelys (Reptilia, Testudinidae). Molecular Ecology, 12, pp. 1415–1424. https://doi.org/10.1046/j.1365-294x.2003.01842.x

Balmer, O., Ciofi, C., Galbraith, D. A., Swingland, I. R., Zug, G. R., & Caccone, A. (2011). Population genetic structure of Aldabra giant tortoises. The Journal of Heredity, 102(1), 29–37.

Bay, R. A., Taylor, E. B., & Schluter, D. (2019). Parallel introgression and selection on introduced alleles in a native species. Molecular Ecology, 28, 2802–2813.

Bolger, A. M., Lohse, M., & Usadel, B. (2014). Trimmomatic: a flexible trimmer for Illumina sequence data. Bioinformatics, 30(15), 2114–2120.

Bourn, D., & Coe, M. (1978). The size, structure and distribution of the giant tortoise population of Aldabra. Philosophical Transactions of the Royal Society of London. B, Biological Sciences, 282(988), 139–175.

Bourn, D., Gibson, C., Augeri, D., Wilson, C. J., Church, J., & Hay, S. I. (1999). The rise and fall of the Aldabran giant tortoise population. Proceedings of the Royal Society B: Biological Sciences, 266(1424), 1091–1100.

Breusing, C., Johnson, S. B., Vrijenhoek, R. C., & Young, C. R. (2019). Host hybridization as a potential mechanism of lateral symbiont transfer in deep-sea vesicomyid clams. Molecular Ecology, 28(21), 4697–4708.

Bunbury, N., von Brandis, R., Currie, J. C., van de Crommenacker, J., Accouche, W., Birch, D., Chong-Seng, L., Doak, N., Haupt, P., Haverson, P., Jean-Baptiste, M., Fleischer-Dogley, F. (2018). Late stage dynamics of a successful feral goat eradication from the UNESCO World Heritage site of Aldabra Atoll, Seychelles. Biological Invasions, 20(7), 1735–1747.

Calsbeek, R., & Smith, T. B. (2003). Ocean currents mediate evolution in island lizards. Nature, 426(6966), 552–555.

Çilingir, F. G., Hansen, D., Ozgul, A., & Grossen, C. (2021). Design of SNP markers for Aldabra giant tortoises using low coverage ddRAD-seq. Conservation Genetics Resources. https://doi.org/10.1007/s12686-021-01225-4

Corlett, R. T. (2017). A bigger toolbox: biotechnology in biodiversity conservation. Trends in Biotechnology, 35(1), 55–65.

Cristofari, R., Plaza, P., Fernández, C. E., Trucchi, E., Gouin, N., Le Bohec, C., Zavalga, C., Alfaro-Shigueto, J., Luna-Jorquera, J & G. (2019). Unexpected population fragmentation in an endangered seabird: the case of the Peruvian diving-petrel. Scientific Reports, 9(1), 2021.

Davey, J. W., & Blaxter, M. L. (2010). RADSeq: next-generation population genetics. Briefings in Functional Genomics, 9(5-6), 416–423.

Davey, J. W., Cezard, T., Fuentes-Utrilla, P., Eland, C., Gharbi, K., & Blaxter, M. L. (2013). Special features of RAD Sequencing data: implications for genotyping. Molecular Ecology, 22(11), 3151–3164.

David, F. N., & Tukey, J. W. (1977). Exploratory data analysis. Biometrics, 33, p. 768. https://doi.org/10.2307/2529486

Fox, E. A., Wright, A. E., Fumagalli, M., & Vieira, F. G. (2019). ngsLD: evaluating linkage disequilibrium using genotype likelihoods. Bioinformatics, 35(19), 3855–3856.

Frankham, R., Ballou, S. E. J., Briscoe, D. A., & Ballou, J. D. (2007). Introduction to Conservation Genetics. Cambridge University Press.

Fumagalli, M., Vieira, F. G., Linderoth, T., & Nielsen, R. (2014). ngsTools: methods for population genetics analyses from next-generation sequencing data. Bioinformatics, 30(10), 1486–1487.

Gibson, C. W. D., & Hamilton, J. (1984). Population processes in a large herbivorous reptile: the giant tortoise of Aldabra Atoll. Oecologia, 61(2), 230–240.

Griffiths, C. J., Hansen, D. M., Jones, C. G., Zuël, N., & Harris, S. (2011). Resurrecting extinct interactions with extant substitutes. Current Biology, 21(9), 762–765.

Griffiths, C. J., Zuël, N., Jones, C. G., Ahamud, Z., & Harris, S. (2013). Assessing the potential to restore historic grazing ecosystems with tortoise ecological replacements. Conservation Biology, 27(4), 690–700.

Grubb, P. (1971). The growth, ecology and population structure of giant tortoises on Aldabra. Philosophical Transactions of the Royal Society B, 260(836), 327–372.

Hansen, D. M. (2015). Non-native megaherbivores: the case for novel function to manage plant invasions on islands. AoB Plants, 7, lv085.

Hansen, D. M., Austin, J. J., Baxter, R. H., de Boer, E. J., Falcón, W., Norder, S. J., Rijsdijk, K. F., Thébaud, C., Bunbury, N., Warren, B. H. (2017). Origins of endemic island tortoises in the western Indian Ocean: a critique of the human-translocation hypothesis. Journal of Biogeography, Vol. 44, pp. 1430–1435. https://doi.org/10.1111/jbi.12893

Hansen, D. M., Donlan, C. J., Griffiths, C. J., & Campbell, K. J. (2010). Ecological history and latent conservation potential: large and giant tortoises as a model for taxon substitutions. Ecography, 33, 272–284.

Hansen, D. M., Josh Donlan, C., Griffiths, C. J., & Campbell, K. J. (2010). Ecological history and latent conservation potential: large and giant tortoises as a model for taxon substitutions. Ecography, 33, 272–284.

Hardy, G. H. (1908). Mendelian proportions in a mixed population. Science, 28(706), 49–50.

Hawlitschek, O., Ramírez Garrido, S., & Glaw, F. (2017). How marine currents influenced the widespread natural overseas dispersal of reptiles in the Western Indian Ocean region. Journal of Biogeography, 44(6), 1435–1440.

Hayden, E. C. (2014). Technology: The $1,000 genome. Nature, 507(7492), 294–295.

Hnatiuk, R. J., Woodell, S. R. J., & Bourn, D. M. (1976). Giant tortoise and vegetation interactions on Aldabra atoll—part 2: coastal. Biological Conservation, 9(4), 305–316.

Hoban, S., Bruford, M. W., Chris Funk, W., Galbusera, P., Patrick Griffith, M., Grueber, C. E., Heuertz, M., Hunter, M. E., Hvilsom, C., Stroil, B. K., Kershaw, F., Khoury, C. K., Laikre, L., Lopes-Fernandes, M., MacDonald, A. J., Mergeay, J., Meek, M., Mittan, C., Mukassabi, T. A., O’Brien, D., Ogden, R., Palma-Silva, C., Ramakrishnan, U., Segelbacher, G., Shaw, R. E., Sjögren-Gulve, P., Veličković, N., Vernesi, C. (2021). Global commitments to conserving and monitoring genetic diversity are now necessary and feasible. BioScience, 71(9), 964–976.

Hohenlohe, P. A., Amish, S. J., Catchen, J. M., Allendorf, F. W., & Luikart, G. (2011). Next-generation RAD sequencing identifies thousands of SNPs for assessing hybridization between rainbow and westslope cutthroat trout. Molecular Ecology Resources, 11, 117–122.

Kehlmaier, C., Graciá, E., Campbell, P. D., Hofmeyr, M. D., Schweiger, S., Martínez-Silvestre, A., Joyce, W., Fritz, U. (2019). Ancient mitogenomics clarifies radiation of extinct Mascarene giant tortoises (Cylindraspis spp.). Scientific Reports, 9(1), 17487.

Kersten, O., Star, B., Leigh, D. M., Anker-Nilssen, T., Strøm, H., Danielsen, H. S. J., Descamps, S., Erikstad, K. E., Fitzsimmons, M. G., Fort, J., Hansen, E. J., Harris, M. P., Irestedt, M., Kleven, O., Mallory, M. L., Jakobsen, K. S., Boessenkool, S. (2021). Complex population structure of the Atlantic puffin revealed by whole genome analyses. Communications Biology, 4(1), 922.

Kopelman, N. M., Mayzel, J., Jakobsson, M., Rosenberg, N. A., & Mayrose, I. (2015). Clumpak: a program for identifying clustering modes and packaging population structure inferences across K. Molecular Ecology Resources, 15(5), 1179–1191.

Korneliussen, T. S., Albrechtsen, A., & Nielsen, R. (2014). ANGSD: analysis of next generation sequencing data. BMC Bioinformatics, 15, 356.

Lepais, O., & Weir, J. T. (2014). SimRAD: an R package for simulation-based prediction of the number of loci expected in RADseq and similar genotyping by sequencing approaches. Molecular Ecology Resources, 14(6), 1314–1321.

Li, H. (2011). Improving SNP discovery by base alignment quality. Bioinformatics, 27(8), 1157–1158.

Li, H., & Durbin, R. (2009). Fast and accurate short read alignment with Burrows-Wheeler transform. Bioinformatics, 25(14), 1754–1760.

Li, H., Handsaker, B., Wysoker, A., Fennell, T., Ruan, J., Homer, N., Marth, G., Abecasis, G., Durbin, R., 1000 Genome Project Data Processing Subgroup. (2009). The sequence alignment/map format and SAMtools. Bioinformatics, 25(16), 2078–2079.

Lou, R. N., Jacobs, A., Wilder, A., & Therkildsen, N. O. (2021). A beginner’s guide to low-coverage whole genome sequencing for population genomics. Molecular Ecology, 00, 1–28 https://doi.org/10.1111/mec.16077

McKenna, A., Hanna, M., Banks, E., Sivachenko, A., Cibulskis, K., Kernytsky, A.,Garimella, K., Altshuler, D., Gabriel, S., Daly, M., DePristo, M. A. (2010). The genome analysis toolkit: a MapReduce framework for analyzing next-generation DNA sequencing data. Genome Research, 20(9), 1297–1303.

Meisner, J., & Albrechtsen, A. (2018). Inferring population structure and admixture proportions in low-depth NGS data. Genetics, 210(2), 719–731.

Merton, L. F. H., Bourn, D. M., & Hnatiuk, R. J. (1976). Giant tortoise and vegetation interactions on aldabra atoll—part 1: inland. Biological Conservation, 9(4), 293–304.

O’Leary, S. J., Puritz, J. B., Willis, S. C., Hollenbeck, C. M., & Portnoy, D. S. (2018). These aren’t the loci you’re looking for: Principles of effective SNP filtering for molecular ecologists. Molecular Ecology. https://doi.org/10.1111/mec.14792

Palkovacs, E. P., Gerlach, J., & Caccone, A. (2002). The evolutionary origin of Indian Ocean tortoises (Dipsochelys). Molecular Phylogenetics and Evolution, 24(2), 216–227.

Pasaniuc, B., Rohland, N., McLaren, P. J., Garimella, K., Zaitlen, N., Li, H., Gupta N., NEale, B. M., Daly, M. J., Sklar, P., Sullivan, P. F., Bergen, S., Moran, J. L., Hultman, C. M., Lichtenstein, P., Magnusson, P., Purcell, S. M., Haas, D. W., Liang. L., Sunyaev, S., Patterson, N., de Bakker, P. I. W., Reich, D., Price, A. L. (2012). Extremely low-coverage sequencing and imputation increases power for genome-wide association studies. Nature Genetics, 44(6), 631–635.

Peart, C. R., Tusso, S., Pophaly, S. D., Botero-Castro, F., Wu, C.-C., Aurioles-Gamboa, D., Baird, A. B., Bickham, J. W., Fircada, J., Galimberti, F., Gemmell, N. J., Hoffman, J. I., Kovacs, K. M., Kunnasranta, M., Lydersen, C., Nyman, T., de Oliveira, L. R., Orr, A. J., Sanvito, S., Valtonen, M., Shafer, A. B. A., Wolf, J. B. W. (2020). Determinants of genetic variation across eco-evolutionary scales in pinnipeds. Nature Ecology & Evolution, 4(8), 1095–1104.

Pedrono, M., Griffiths, O. L., Clausen, A., Smith, L. L., Griffiths, C. J., Wilmé, L., & Burney, D. A. (2013). Using a surviving lineage of Madagascar’s vanished megafauna for ecological restoration. Biological Conservation, Vol. 159, pp. 501–506. https://doi.org/10.1016/j.biocon.2012.11.027

Peterson, B. K., Weber, J. N., Kay, E. H., Fisher, H. S., & Hoekstra, H. E. (2012). Double digest RADseq: an inexpensive method for de novo SNP discovery and genotyping in model and non-model species. PloS ONE, 7(5), e37135.

Pritchard, J. K., Stephens, M., & Donnelly, P. (2000). Inference of population structure using multilocus genotype data. Genetics, 155(2), 945–959.

Quesada, V., Freitas-Rodríguez, S., Miller, J., Pérez-Silva, J. G., Jiang, Z.-F., Tapia, W., Santiago-Fernández, O., Campos-Iglesias, D., Kuderna, L. F. K., Quinzin, M., Álvarez, M., G., Carrero, D., Beheregaray, L. B., Gibbs, J. B., Chiari, Y., Glaberman, S., Ciofi, C., Araujo-Voces, M., Mayoral, P., Arango, J. R., Tamargo-Gómez, I., Roiz-Valle, D., Pascual-Torner, M., Evans, B. R., Edwards, D. L., Garrick, R. C., Russello, M. A., Poulakakis, N., Gaughran, S. J., Rueda, D. O., Bretines, G., Marquès-Bonet, T., White, K. P., Caccone, A., López-Otín, C. (2019). Giant tortoise genomes provide insights into longevity and age-related disease. Nature Ecology & Evolution, 3(1), 87–95.

Quinlan, A. R., & Hall, I. M. (2010). BEDTools: a flexible suite of utilities for comparing genomic features. Bioinformatics, 26(6), 841–842.

R Core Team. (2020). R: A language and environment for statistical computing (Version R Foundation for Statistical Computing, Vienna, Austria).

Reed, D. H. (2005). Relationship between population size and fitness. Conservation Biology, 19(2), 563–568.

Reed, D. H., & Frankham, R. (2003). Correlation between fitness and genetic diversity. Conservation Biology, 17(1), 230–237.

Rochette, N. C., Rivera-Colón, A. G., & Catchen, J. M. (2019). Stacks 2: Analytical methods for paired-end sequencing improve RADseq-based population genomics. Molecular Ecology, 28, 4737–4754.

Shafer, A. B. A., Wolf, J. B. W., Alves, P. C., Bergström, L., Bruford, M. W., Brännström, I., Bruford, M. W., Brännström, I., Colling, G., Dalén, L., Meester, L. D., Ekblom, R., Fawcett, K. D., Fior, S., Hajibabaei, M., Hill, J. A., Hoezel, A. R., Höglund, J., Jensen, E. L., Krause, J., Kristensen, T. N., Krützen, M., McKay, J. K., Norman, A. J., Ogden, R., Österling, E. M., Ouborg, N. J., Piccolo, J., Popović, D., Primmer, C. R., Reed, F. A., Roumet, M., Salmona, J., Schenekar, T., Schwartz, M. K., Segelbacher, G., Senn, H., Thaulow, J., Valtonen, M., Veale, A., Vergeer, P., Vijay, N., Vilà, C., Weissensteiner, M., Wennerström, L., Wheat, W., Zieliński, P. (2015). Genomics and the challenging translation into conservation practice. Trends in Ecology & Evolution, 30(2), 78–87.

Silva, P., López-Bao, J. V., Llaneza, L., Álvares, F., Lopes, S., Blanco, J. C., Cortés, Y., García, E., Palacios, V., Rio-Maior, H., Ferrand, N., Godinho, R. (2018). Cryptic population structure reveals low dispersal in Iberian wolves. Scientific Reports, 8(1), 14108.

Skotte, L., Korneliussen, T. S., & Albrechtsen, A. (2013). Estimating individual admixture proportions from next generation sequencing data. Genetics, 195(3), 693–702.

Stoddart, D. R., & Peake, J. F. (1979). Historical records of Indian Ocean giant tortoise populations. Philosophical Transactions of the Royal Society of London. Series B, Biological Sciences, 286, 147–161.

Swingland, I. R., North, P. M., Dennis, A., & Parker, M. J. (1989). Movement patterns and morphometrics in giant tortoises. The Journal of Animal Ecology, 58(3), 971–985.

Tajima, F. (1989). Statistical method for testing the neutral mutation hypothesis by DNA polymorphism. Genetics, 123(3), 585–595.

Turnbull, L. A., Ozgul, A., Accouche, W., Baxter, R., ChongSeng, L., Currie, J. C., Doak, N., Hansen, D. M., Pistorius, P., Richards, H., van de Crommenacker, J., von Brandis, R., Fleischer-Dogley, F., Bunbury, N. (2015). Persistence of distinctive morphotypes in the native range of the CITES-listed Aldabra giant tortoise. Ecology and Evolution, 5(23), 5499–5508.

Turtle Taxonomy Working Group [Rhodin, A., G. J., Iverson, B., J., Bour, R., Fritz, U., Georges, A., Shaffer, B., H., Dijk, V., &] P. P. (2017). Turtles of the World: Annotated Checklist and Atlas of Taxonomy, Synonymy, Distribution, and Conservation Status (8th Ed.): Conservation Biology of Freshwater Turtles and Tortoises: a Compilation Project of the IUCN/SSC Tortoise and Freshwater Turtle Specialist Group.

Walton, R., Baxter, R., Bunbury, N., Hansen, D., Fleischer-Dogley, F., Greenwood, S., & Schaepman-Strub, G. (2019). In the land of giants: habitat use and selection of the Aldabra giant tortoise on Aldabra Atoll. Biodiversity and Conservation, 28, 3183–3198.

Warmuth, V. M., & Ellegren, H. (2019). Genotype-free estimation of allele frequencies reduces bias and improves demographic inference from RADSeq data. Molecular Ecology Resources, 19(3), 586–596.

Watterson, G. A. (1975). On the number of segregating sites in genetical models without recombination. Theoretical Population Biology, 7(2), 256–276.

White, C., Selkoe, K. A., Watson, J., Siegel, D. A., Zacherl, D. C., & Toonen, R. J. (2010). Ocean currents help explain population genetic structure. Proceedings of the Royal Society B: Biological Sciences, 277(1688), 1685–1694.

Wickham, H. (2016). ggplot2: elegant graphics for data analysis. Retrieved from https://ggplot2.tidyverse.org

Wietlisbach, X. (2017). The genetics of giants: how have time and space shaped genetic variation within and among Aldabra giant tortoise populations? (MSc.; E. Postma, Ed.). University of Zurich.

Záveská, E., Maylandt, C., Paun, O., Bertel, C., Frajman, B., The Steppe Consortium, & Schönswetter, P. (2019). Multiple auto- and allopolyploidisations marked the Pleistocene history of the widespread Eurasian steppe plant *Astragalus onobrychis* (Fabaceae). Molecular Phylogenetics and Evolution, 139, 106572.

