## Supplementary material for "Low-coverage reduced representation sequencing reveals subtle within-island genetic structure in Aldabra giant tortoises": Suppl. Table 2: Suppl. Table 2.docx

**Suppl. Table 2** Pairwise Fst estimations with the downsampled dataset

|  | **Malabar Group 1** | **Malabar Group 2** | **Grande Terre East** | **Grande Terre West** | **Picard** |
| --- | --- | --- | --- | --- | --- |
| **Malabar Group 1** | 0 |  |  |  |  |
| **Malabar Group 2** | 0.036 | 0 |  |  |  |
| **Grande Terre East** | 0.055 | 0.039 | 0 |  |  |
| **Grande Terre West** | 0.012 | 0.024 | 0.026 | 0 |  |
| **Picard** | 0.005 | 0.006 | 0.017 | 0 | 0 |
