## Supplementary material for "Low-coverage reduced representation sequencing reveals subtle within-island genetic structure in Aldabra giant tortoises": Suppl. Fig. 1-6: Suppl. Figures 1-6.docx

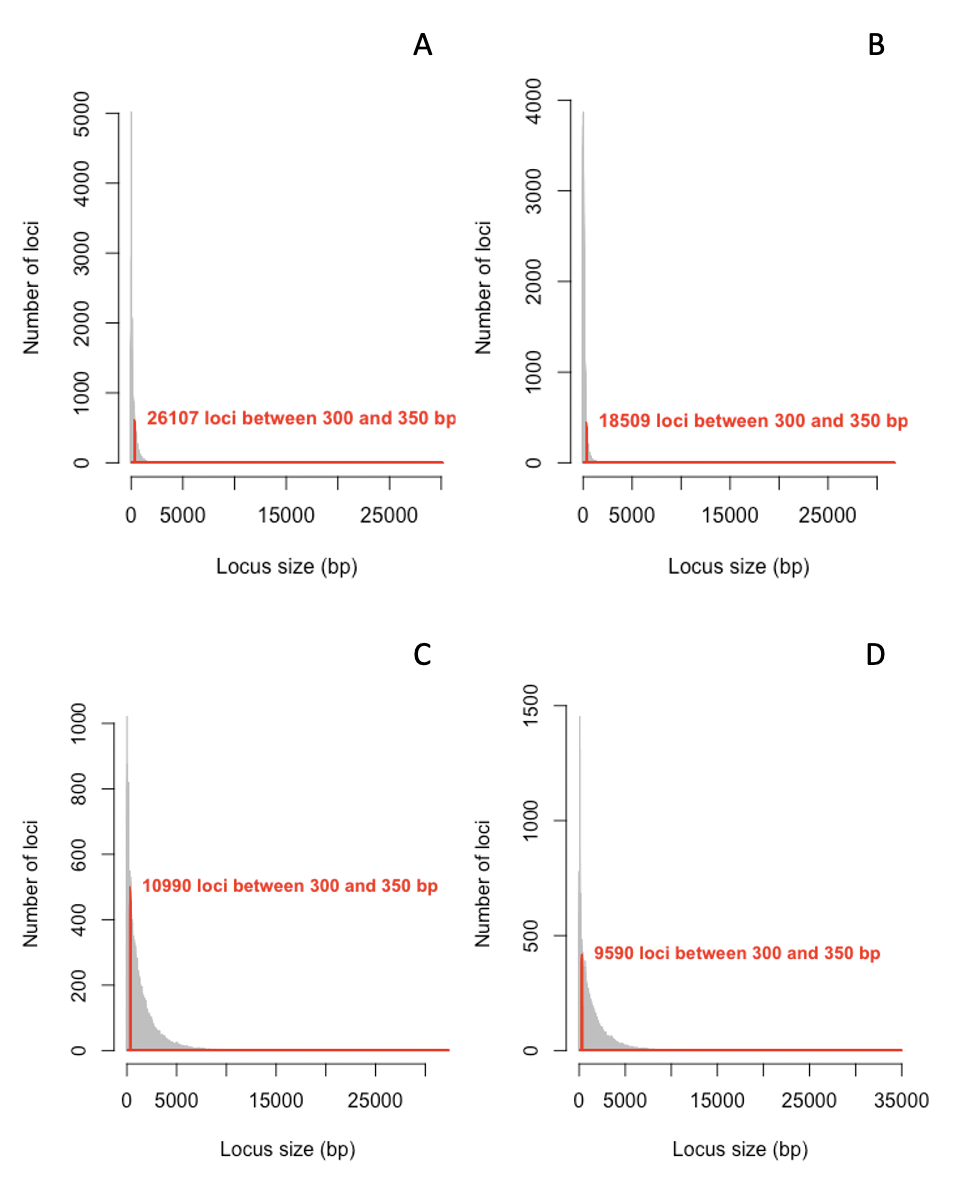

**Suppl. Fig. 1** *In silico* double digests of the 50% of the *C. abingdonii* genome with A) EcoRI and BfaII B) EcoRI and MseI, C) EcoRI and MspI, and D) EcoRI and TaqI with the selection window of 350-400bp is shown.

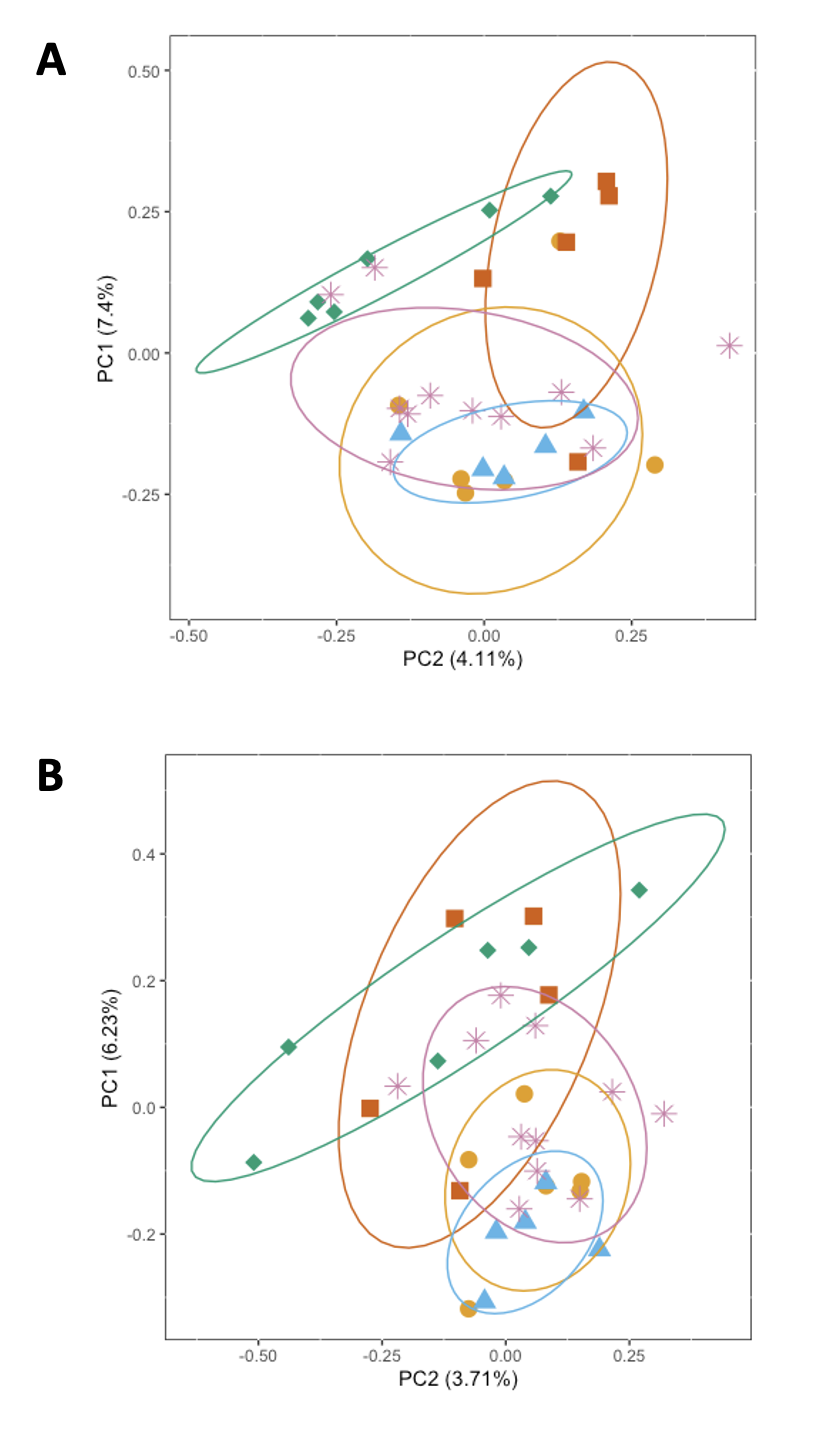

**Suppl. Fig. 2** PCA plot **A)** generated with 6,153 SNPs with MAF > 0.01 from the main dataset. PC1 represents 7.4% of the total variance whereas PC2 represents 4.11% of it, **B)** with 1,632 SNPs with MAF > 0.01 from the downsampled dataset. PC1 represents 6.23% of the total variance whereas PC2 represents 3.71% of it. For both of the figures blue triangles show individuals from Malabar Group 1, green diamonds show from Malabar Group 2, red squares show from Grande Terre East, orange circles show from Grande Terre West and pink asterisks show individuals from Picard.

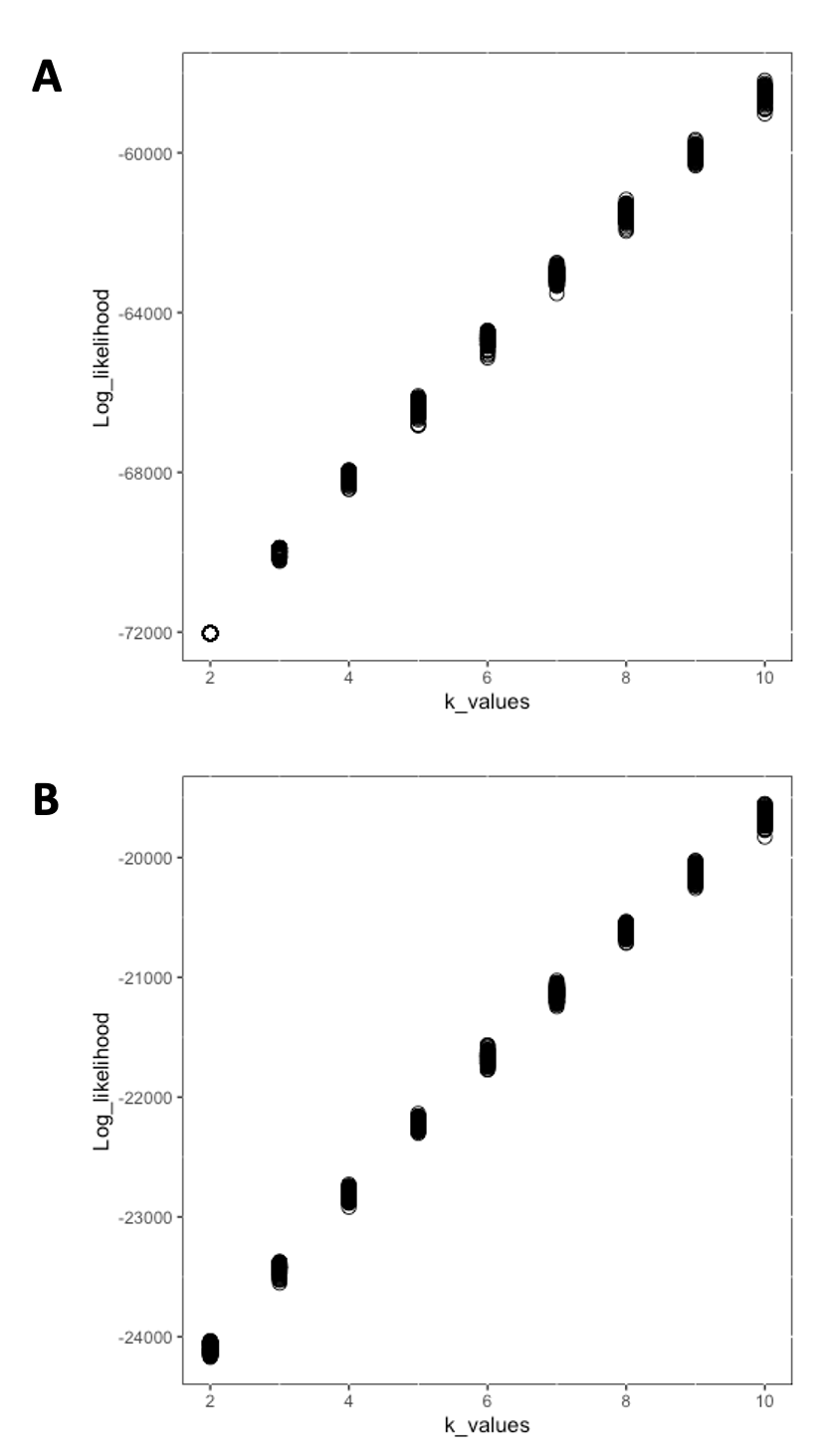

**Suppl. Fig 3** Log likelihood values of 100 ngsAdmix runs per k=2-10 are shown. Panel **A** and **B** represent the main and the downsampled datasets, respectively.

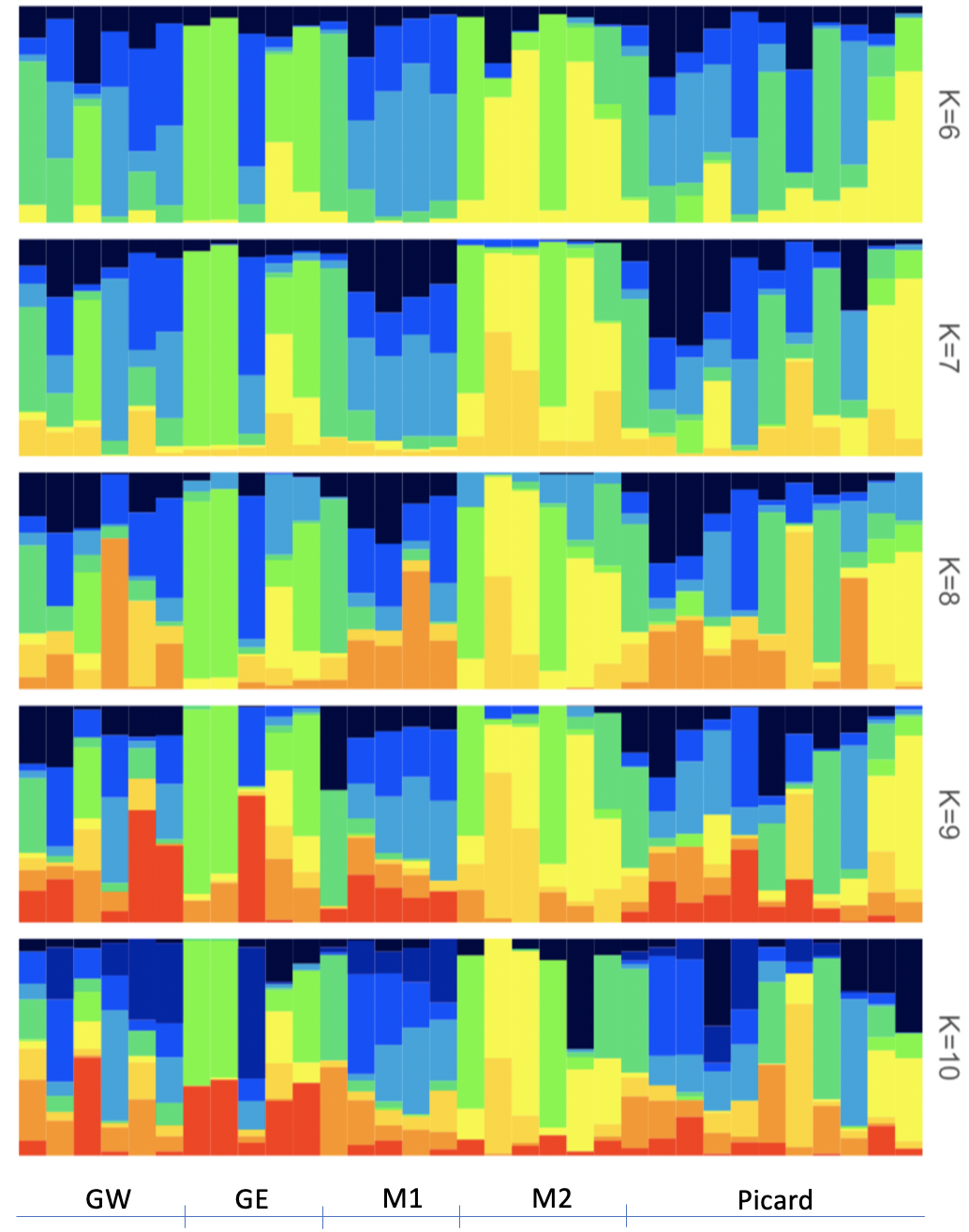

**Suppl. Fig. 4** Results of ngsAdmix runs performed with the main dataset at k=5-10 are shown. GW, GE, M1, and M2 stand for Grande Terre West, Grande Terre East, Malabar Group 1, and Malabar Group 2, respectively.

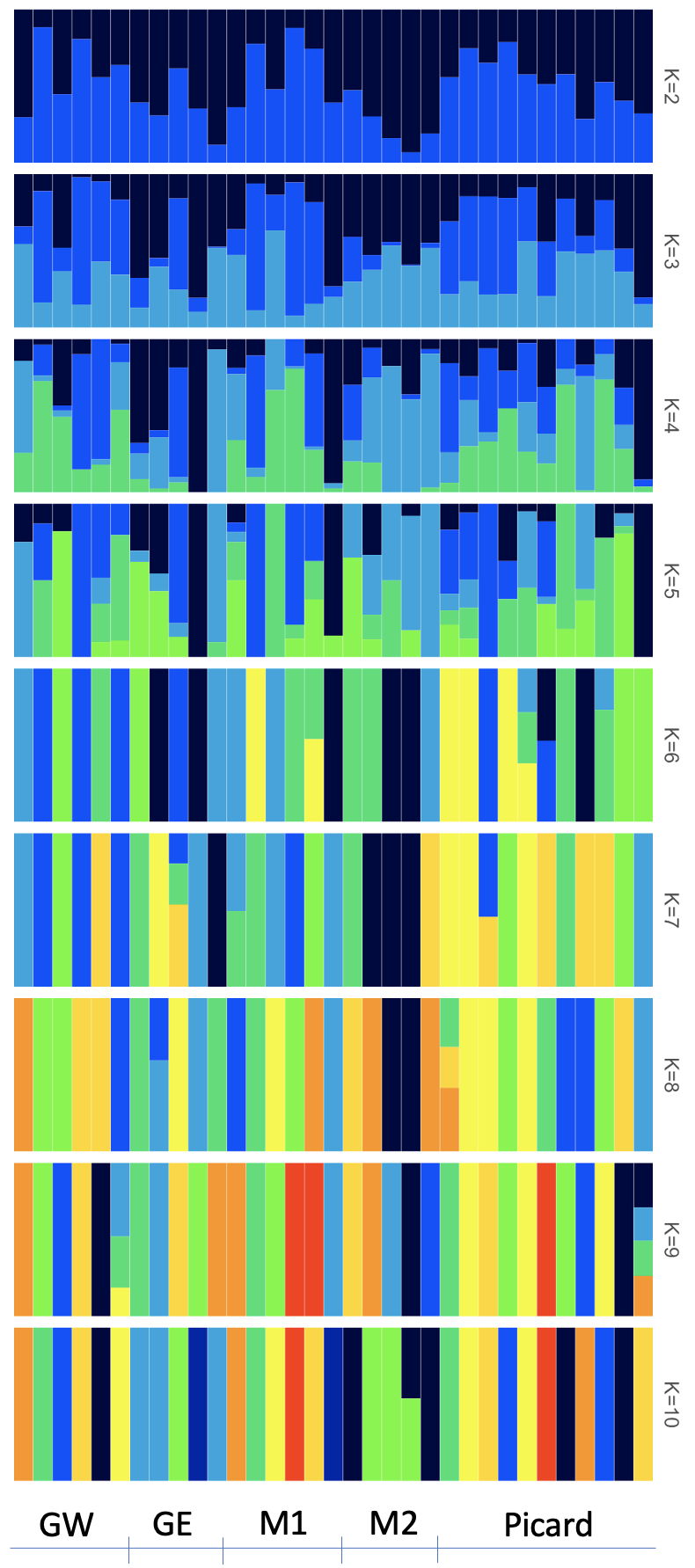

**Suppl. Fig. 5** Results of ngsAdmix runs performed with the downsampled dataset at k=2-10 are shown. GW, GE, M1, and M2 stand for Grande Terre West, Grande Terre East, Malabar Group 1 and Malabar Group 2, respectively.

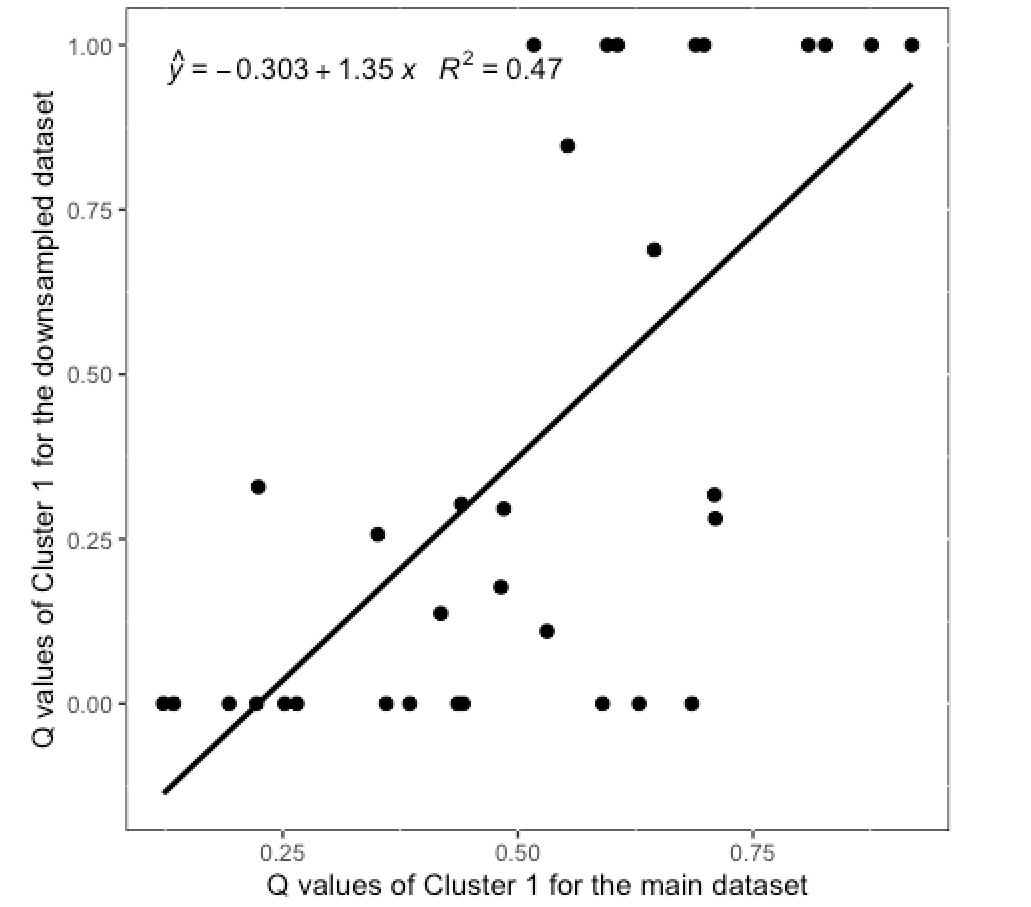

**Suppl. Fig. 6** Linear regression between the admixture proportions (q values) of the Cluster 1 of the main and the downsampled datasets are shown.
